# iPSC-Astrocyte morphology reflects patient clinical markers

**DOI:** 10.1101/2023.07.15.548687

**Authors:** Helen A. Rowland, Georgina Miller, Qiang Liu, Nicola Sharp, Bryan Ng, Tina Wei, Kanisa Arunasalam, Ivan Koychev, Anne Hedegaard, Elena M. Ribe, Dennis Chan, Tharani Chessell, Ece Kocagoncu, Jennifer Lawson, Paresh Malhotra, Basil H. Ridha, James B. Rowe, Alan J. Thomas, Giovanna Zamboni, Henrik Zetterberg, M. Zameel Cader, Richard Wade-Martins, Simon Lovestone, Alejo Nevado-Holgado, Andrey Kormilitzin, Noel J. Buckley

## Abstract

Human iPSCs provide powerful cellular models of Alzheimer’s disease (AD) and offer many advantages over non-human models, including the potential to reflect variation in individual-specific pathophysiology and clinical symptoms Previous studies have demonstrated that iPSC-neurons from individuals with Alzheimer’s disease (AD) reflect clinical markers, including β-amyloid (Aβ) levels and synaptic vulnerability. However, despite neuronal loss being a key hallmark of AD pathology, many risk genes are predominantly expressed in glia, highlighting them as potential therapeutic targets. In this work iPSC-derived astrocytes were generated from a cohort of individuals with high versus low levels of the inflammatory marker YKL-40, in their cerebrospinal fluid (CSF). iPSC-derived astrocytes were treated with exogenous Aβ oligomers and high content imaging demonstrated a correlation between astrocytes that underwent the greatest morphology change from patients with low levels of CSF-YKL-40 and more protective *APOE* genotypes. This finding was subsequently verified using similarity learning as an unbiased approach. This study shows that iPSC-derived astrocytes from AD patients reflect key aspects of the pathophysiological phenotype of those same patients, thereby offering a novel means of modelling AD, stratifying AD patients and conducting therapeutic screens.

**Graphical Abstract:** 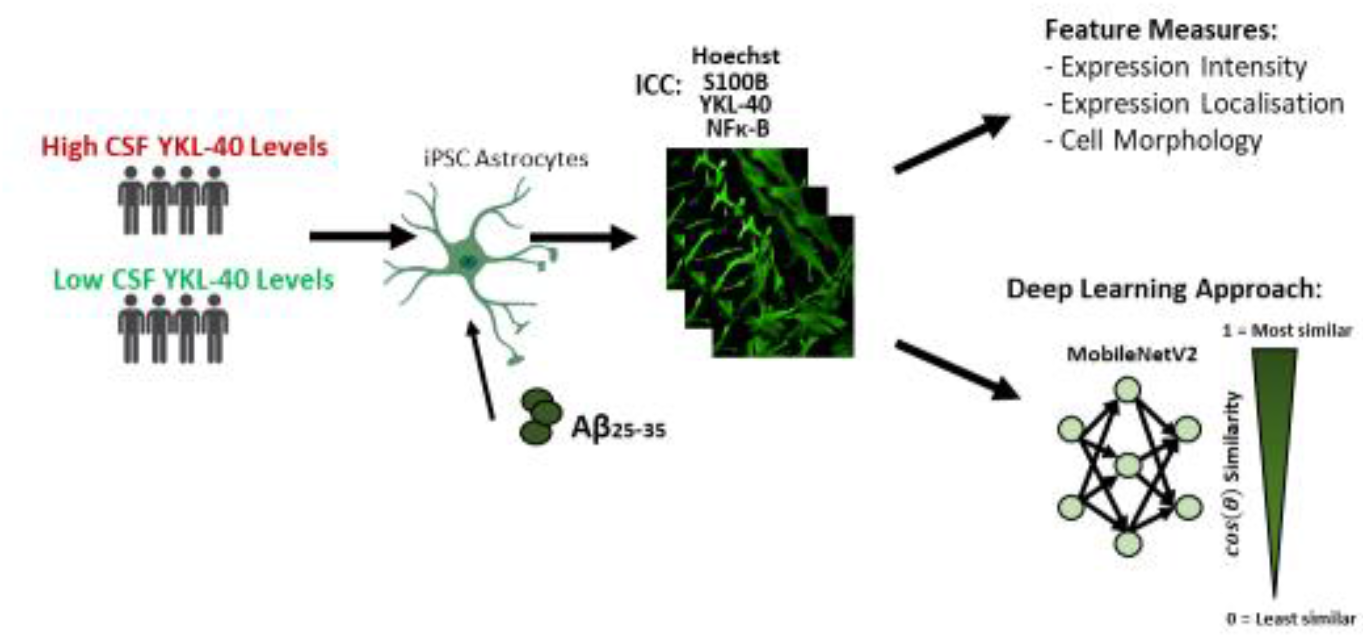

## Introduction

Human iPSCs are increasingly used to model Alzheimer’s disease, whereas many non-human *in vitro* and *in vivo* models have failed to recapitulate key aspects of disease phenotype. The majority of studies utilising stem cells from patients have focused on disease variants in the causative genes *APP, PSEN1* or *PSEN2*. Studies using sporadic AD (sAD) iPSC lines have not always exhibited clear disease phenotypes [1, 2]. However, recent studies have demonstrated that clinical markers are indeed reflected by iPSC-derived neurons from individuals with sAD, including secreted β-amyloid (Aβ) levels and synaptic vulnerability [3, 4]. These findings open the feasibility of modelling sAD, allowing the interrogation of mechanisms and identification of therapeutic targets in a personalised medicine approach.

Despite neuronal loss being a key hallmark of AD pathology, GWAS studies have identified many sAD risk genes that are predominantly expressed in glia, including the largest sAD risk gene, *APOE*, highlighting them as important potential therapeutic targets [5]. Astrocytes play a role in many physiological functions in the brain, but their diversity and role in disease pathogenesis is not well characterised, in particular their contribution to disease under reactive states. With the capacity to generate reactive astrocytes from patients, these questions can now be addressed [6].

The role of inflammation in AD, and the contribution of glial cells to this, is widely recognised and is being increasingly explored as a therapeutic target. Currently several markers of inflammation including TNFα, IL-6, CRP, and YKL-40 have been identified as important signalling molecules that affect the brain of patients with dementia. CSF YKL-40 is increased in the prodromal and preclinical phase of AD, making it an important biomarker of inflammation in AD [7]. How YKL-40 is affected in AD on cellular level in glial cells has yet to be determined. To this end, we have conducted a preliminary study to determine if iPSC-derived astrocytes from patients of the Deep and Frequent Phenotyping (DFP) study cohort [6], demonstrating astrocyte changes in morphology in response to Aβ burden, reflect the differences in *APOE* genotype and CSF YKL-40 levels of the donor patients.

## Methods

### Generation and Maintenance of iPSCs and astrocytes from the DFP Cohort

The Deep and Frequent Phenotyping (DFP) cohort pilot study protocol was previously published [8] and the generation of 14 iPSC lines from a subset of early symptomatic Alzheimer’s disease cases also described [4]. 8 of these lines were selected representing the 4 highest and 4 lowest patient CSF YKL-40 levels. iPSCs were maintained on Geltrex (Gibco) matrix protein coated plates and cultured in mTeSR^™^ medium (STEMCELL tech), passaged using Versene (Gibco) as cells reached 80% confluence. All cells were maintained at 37°C, 5% CO_2_, and 20% O_2_. Differentiation of iPSCs to astrocytes was done by modifying a previously described protocol [9]; *details are in supplementary data*.

The collection of clinical data from the DFP Cohort has also been described [4, 8]. CSF YKL-40 concentration was measured using a commercially available immunoassay according to the manufacturer’s instructions (MesoScale Discovery, Rockville, MD). *APOE* status for each of the patients had previously been determined **(Supplementary Figure 1)**, and for this study when compared to other clinical measures and cell data, were converted into a numerical scale for all genotypes dependent on risk where *APOE* ε2/ε3 would have a value of 1 and ε4/ε4 a value of 4.

### Aβ_25-35_ Oligomer Preparation and Treatment

Aβ_25-35_ was selected as it retains toxicity of the full-length Aβ peptide, and still maintains the essential domain of Aβ_1-42_ aggregation, but its stability is not limited *in vitro* as the Aβ_1-42_ aggregated fibrillary state is [10, 11]. Previous work has demonstrated that synaptic loss in response to extrinsic Aβ peptides was patient specific and reproducible with both Aβ peptides, suggesting that Aβ_25-35_ would be acceptable to explore the effect of Aβ related stress for the purposes of this study [4]. Preparation of Aβ_25-35_ oligomers was carried out as previously described [12]. Aliquots of 2mg/ml were defrosted on ice and diluted in NMM to a final concentration of 10μM. Astrocytes were treated with Aβ for 24 hours.

### Immunocytochemistry

Cells were fixed with 4% paraformaldehyde and stored at 4°C until blocked with 10% donkey serum in Phosphate-Buffered Saline (PBS) (Invitrogen) and 0.25% Triton X-100 (Sigma-Aldrich). After 1 hour’s incubation at RT, primary antibodies YKL-40 [human-chitinase-3-like 1] (Goat, AF2599, Bio-Techne), s100B (Mouse, SAB4200671, Sigma) and NF-κB (Rabbit, D14E12 Cell Signalling) were applied 1:500 in PBS with 5% donkey serum and left overnight at 4°C. Cells were washed with PBS and secondary antibodies 1:500 in PBS were added for 2 hours. After washing again with PBS, 4μM Hoechst 33342 in PBS was added for 5 minutes at RT. Cells were washed with PBS and then stored in PBS at 4°C until imaged in the Opera Phenix (PerkinElmer) at 40x magnification and images analysed using a Harmony software to assess protein expression and morphology *(see supplementary)*.

### Deep Learning: Classification Task and Similarity Learning

Astrocyte images from the s100B channel were pre-processed and input into a deep learning model architecture modified from the MobileNetV2 [13]. Transfer learning was previously applied, where the weights were pretrained on the ImageNet dataset [14]. The output classification layer from MobileNetV2 was removed adding max and global average pooling layers to yield output vectors of 1280 values for each input image. Cosine similarity was used to calculate the similarity of images for pairwise control and Aβ treatment conditions across cell lines *(see supplementary)*.

### Statistical Analyses

Quantitative graphs and statistical analyses were carried out on GraphPad Prism 9.5.0. Similar to previous analyses [4] the data was confirmed to have a normal distribution, and correlation of cell data and clinical data was carried out using Pearson’s correlation coefficient. For characterisation experiments assessing protein expression by immunocytochemistry in patient lines in response to Aβ treatment a two-way ANOVA test, with Sidak’s multiple comparisons test was carried out. Statistical significance was held to P < 0.05, where ^*^ P < 0.05, ^**^ P < 0.01, ^***^ P < 0.001 and ^****^ P < 0.0001

## Results

Increasing evidence puts YKL-40 as a potential biomarker, increased in inflammation and in AD, where CSF YKL-40 levels can be predictive of AD progression. Therefore, we generated iPSC-astrocytes from 8 individuals from the DFP cohort selecting 4 patients with the highest and 4 patients with the lowest YKL-40 CSF levels. Astrocytes were differentiated and matured for over 150 days and expressed glial markers (**Figure 1A, Supplementary Figure 1**). It is important to note, that despite the cell lines being differentiated at the same time, there were large baseline differences in expression of glial markers. We also found that baseline expression of YKL-40 mRNA and protein levels were extremely variable between astrocytes from patients (**Supplementary Figure 1**), with no correlation to the patient demographical or clinical data previously collected and described [4, 8].

**Figure 1:**
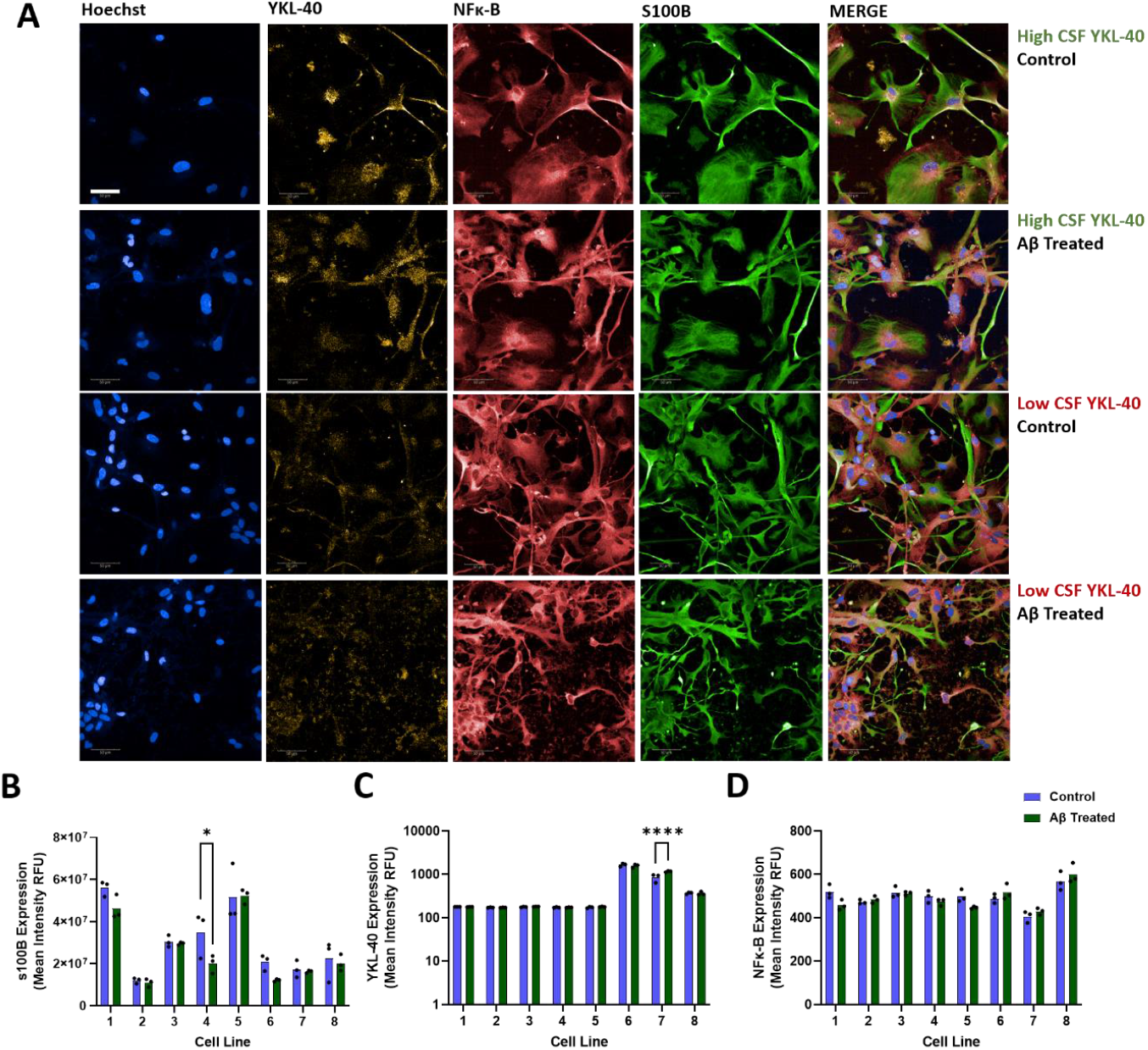
Characterisation of iPSC-derived astrocytes from patients with high and low CSF YKL-40 values. **(A)** Astrocytes were matured to over 150 days where control iPSC-astrocytes (blue) or treated with 10μM Aβ_25-35_ (green) were subsequently stained for Hoechst (blue), YKL-40 (yellow), NF-κB (red), s100B (green). **(B)** Expression of s100B was calculated for each patient line with and without Aβ treatment as determined by mean intensity expression. **(C)** YKL-40 Expression. **(D)** NF-κB Expression. Scale bar represents 50μm.

Previous findings indicate that iPSC neurons vulnerability to Aβ, reflect that of the patients they are derived from [4]. We sought to determine whether iPSC astrocytes also reflected features from their respective patients in response to Aβ. To this end, we treated iPSC astrocytes with Aβ_25-35_ *(see methods)*. We found that treatment with Aβ generally did not alter s100B or YKL-40 protein levels except in a single line (**Figure 1B and 1C**). We also did not detect any changes in protein expression or nuclear localisation of NF-κB (**Figure 1D**).

However we observed that iPSC-astrocytes treated with Aβ underwent significant changes to their morphology. Astrocytes treated with Aβ showed significant process extension in some patients as exampled in **Figure 1A** and quantified in **Figure 2A**. We quantified these morphology changes looking at the ratio of cell width to cell length using the s100B channel and found that this was significantly altered in the patients with low CSF YKL-40 levels. The change in cell width to length and the CSF-YKL-40 levels of the patients produced a strong correlation (**Figure 2B**.) Previous studies have demonstrated a link between CSF YKL-40 in *APOE* ε4 carriers [15]. Here we confirm findings, demonstrating a correlation between higher CSF YKL-40 values and *APOE* ε4 status in the DFP cohort as well (**Figure 2C**). Likewise, we also show a correlation between non-significant changes in cell width to length ratio in response to Aβ treatment and *APOE* status (**Figure 2D**).

**Figure 2:**
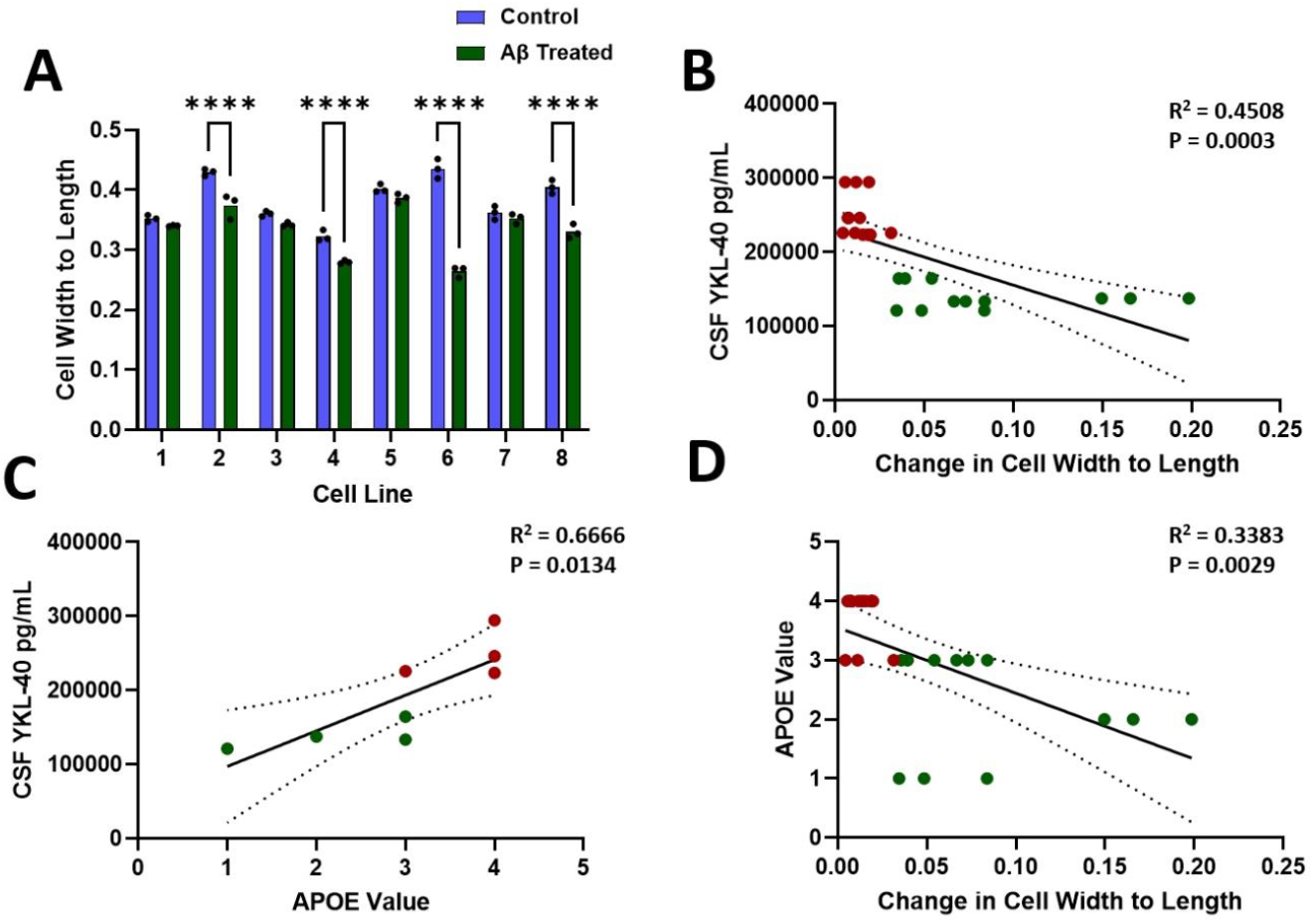
**(A)** Cell Width to Length of control iPSC-astrocytes (blue) or treated with 10μM Aβ_25-35_ (green). **(B)** Correlation of CSF YKL-40 levels to the change in cell width to length between control and Aβ25-35 treated^*^. **(C)** Correlation of CSF YKL-40 to the APOE value of the patient where higher AD risk genotypes equates to a higher APOE value. **(D)** Correlation of APOE value to change in cell width to length between control and Aβ_25-35_ treated. Red represents patients with high levels of CSF YKL-40, with green representing low levels. ^*^*Data remains significant with the removal of patient 7 representing the greatest change in cell width to length (R*^*2*^ *= 0*.*6944, P <0*.*0001)*

This measure of morphology change demonstrates a strong correlation despite a relatively small sample size. However even with best efforts and practice, methods to analyse morphological changes are subject to human bias and software capabilities to accurately segment and capture biologically relevant information. There is potential to lose much information about the numerous changes these astrocytes undergo in response to Aβ treatment. To challenge this, we applied deep learning approaches to take an unbiased view of astrocyte changes between control and Aβ treated conditions from whole field view images. A schematic of this process is shown in **Figure 3A**. The summary of these findings is shown in **Figure 3B** where patients with low CSF YKL-40 values showed less similarity to the control set images. When comparing the change in image similarity of treated and control in each patient from the change of width to length ratio between treatments a significant correlation was seen **(Figure 3C)** indicating that width to length ratio must capture key changes in astrocytes induced by Aβ treatment as a proof of concept. Likewise, this change in image similarity also significantly correlates with the clinical data of patient YKL-40 CSF levels **(Figure 3D)**.

**Figure 3:**
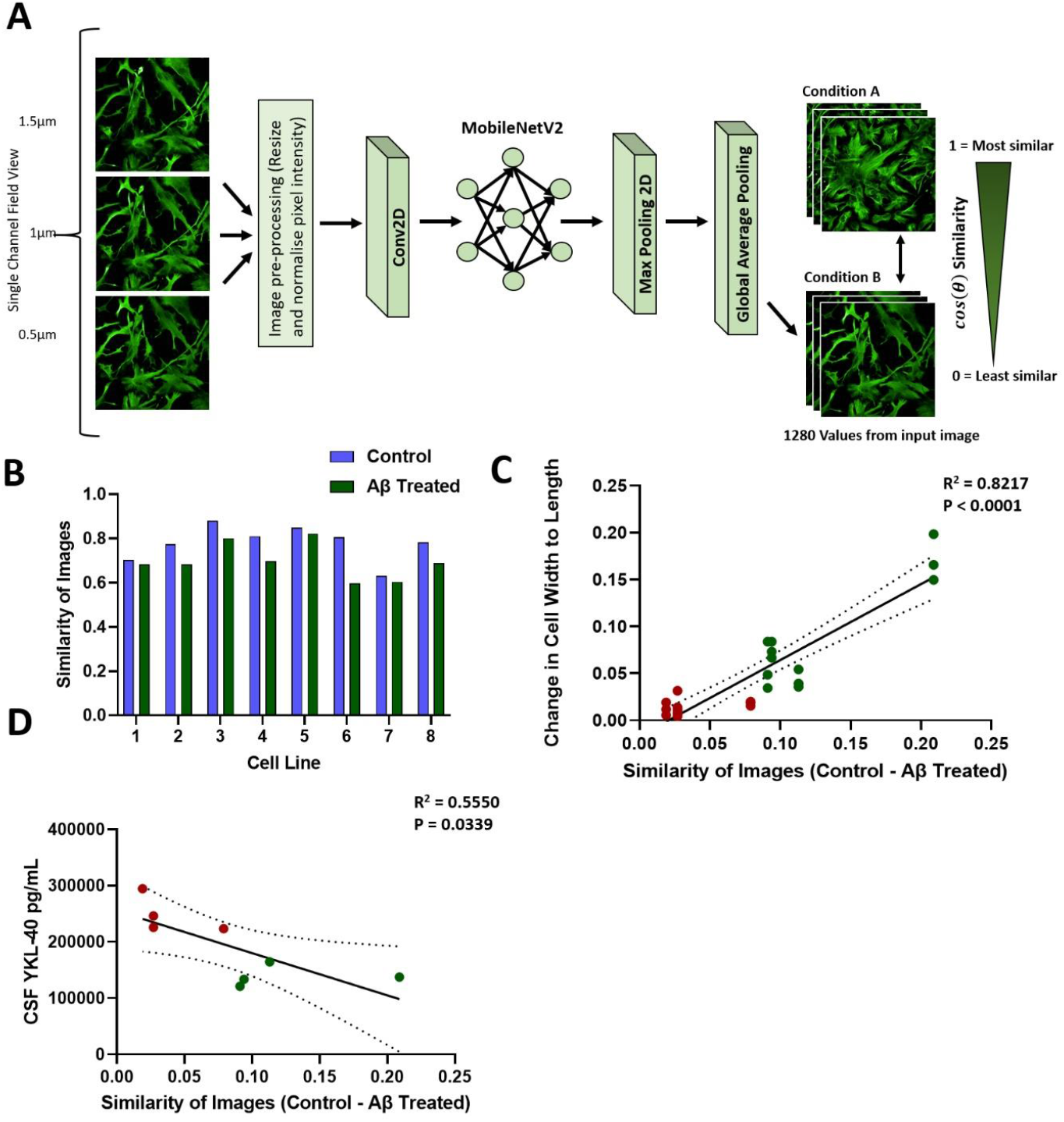
Use of deep learning to compare effect of Aβ treatments in an unbiased approach. **(A)** Schematic demonstrating the network setup using images from the s100B channel. A model was developed to determine how similar images in the s100B channel were from the untreated control of each patient were to its training dataset, and how images of astrocytes treated with Aβ were to their respective controls, Similarity learning was applied where 0 represents no similarity, and 1 an identical image **(B)** Similarity of the images of the test dataset to that of each patient’s control training dataset. The control test dataset is shown in blue, the Aβ treated test dataset is shown in green. **(C)** Correlation of the change in width to length ratio of control and Aβ-treated astrocytes to the similarity of control images minus the similarity of Aβ treated astrocytes (where higher values represent more divergence). **(D)** Correlation of the change in similarity of images to patient YKL-40 CSF levels. Red represents patients with high YKL-40 CSF levels, green represents patients with low YKL-40 CSF levels.

## Discussion

In this work we have shown that astrocyte response to Aβ, as determined by change in morphology after treatment, reflect patient stratification according to CSF YKL-40 levels and *APOE* genotype. This provides further evidence that iPSC cells can be used to model person specific processes that may underlie individual pathophysiology in AD, and extends the evidence from iPSC-derived neurons to iPSC-derived astrocytes as candidates to model patient phenotypes.

The diversity of astrocytes across the CNS is increasingly being appreciated, with steps in advancing our knowledge of various subtypes, inflammatory states, and in disease [16, 17]. Together these varying subtypes, and reactive states result in a large range of morphological diversity that needs to be better characterised and understood. In our own patient cohort we found much morphological diversity within our iPSC-derived astrocytes and this was reflected in large differences in baseline expression of glial markers between lines. This may undermine capacity to directly compare astrocyte phenotypes to patient clinical data at present, but in future studies we hope that greater understanding of causes of morphological changes and diversity in an *in vitro* setting will help standardise methodology and consistency in astrocyte populations that are generated. However, despite this current difficulty in the field, we found that the amount in which astrocytes were morphologically affected by the presence of a toxic stimuli, rather than baseline readouts, strongly correlated with *APOE* genotype and YKL-40 biomarker data despite a relatively small patient cohort. This provides useful evidence of the ways in which astrocytes can be used to model disease or utilised to test therapeutic interventions.

Curiously, astrocytes that exhibited greater morphological change in response to Aβ came from patients with lower CSF YKL-40 levels and more protective *APOE* genotypes. Further studies are needed to understand whether this is a protective or detrimental response. Previous studies have indicated that YKL-40 from plasma is negatively associated with brain Aβ deposition and positively associated with memory performance in people with subjective memory complaint [18]. In this manner astrocytes from patients with high CSF-YKL-40 levels may be acting more protectively. Further understanding of the role of YKL-40 in glial cells will be useful to understand the mechanistic role this protein plays in astrocyte activation and disease pathology. Conversely the growing literature on the role of APOE where *APOE* ε4/ε4 conveys greatest risk in astrocytes have demonstrated impaired function in Aβ clearance, increased inflammation, and metabolic dysregulation among others [19-22]. If astrocyte function is impaired, this may suggest that the early morphological changes in response to Aβ treatment are instead protective. It has yet to be determined how changes in morphology may reflect protective or detrimental effects, and also how different cell types may be differentially affected in AD and if this is accurately reflected in patient clinical data.

In this work we have also exampled, through the use of iPSC-derived astrocytes, a potential tool for therapeutic uses. This proof of concept demonstrates how Deep Learning can be used as an unbiased way to classify cell phenotype, with the clear translational potential for screening of drug libraries. Our data showed that our Deep Learning approach could view the same morphological changes induced by Aβ as conventional means of analysis. By applying this approach, user biases are eliminated as the model will observe global changes rather than a fixed measure, which possibly ignores other salient changes. In using similarity learning, we can attempt to stratify astrocytes from patients and test compounds to see if astrocytes can be ‘restored’ to baseline, or healthy controls. Together the data here highlight that different cell types may respond differently in disease and can be reflected by patient clinical data, adding more evidence towards a personalised medicine approach. In this effort, the use of unbiased approaches that can rapidly stratify and identify hit compounds may make this a more viable option in the future.

## Acknowledgements

This work was funded by a National Institute for Health Research-Medical Research Council Dementias Platform UK Experimental Medicine Award (MR/L023784/2) and an Equipment Award (MR/M024962/1). The Deep and Frequent Phenotyping clinical study is funded by the Medical Research Council (MR/N029941/1). This project was supported by a Stem Cells for Biological Assays of Novel Drugs and Predictive Toxicology funding from the Innovative Medicines Initiative Joint Undertaking under Grant Agreement Number 115439, resources of which are composed of financial contribution from the European Union’s Seventh Framework Programme (FP7/2007e2013) and European Federation of Pharmaceutical Industries and Associations companies in kind contribution. The work was also supported by the National Institute for Health Research Oxford Biomedical Research Centre. H.A.R. was supported by an Alzheimer’s Research UK Thames Valley Network Early Career Researcher Award. B.N. was supported by the National Science Scholarship by the Agency for Science, Technology and Research, Singapore. I.K. received support through the Oxford Health Biomedical Research Centre, the Medical Research Council and the National Institute for Health Research. B.H.R. was supported by the National Institute for Health Research Biomedical Research Centre at University College London Hospitals and P.M. was supported by the National Institute for Health Research Biomedical Research Centre at Imperial College London. H.Z. is a Wallenberg Scholar supported by grants from the Swedish Research Council (#2022-01018 and #2019-02397), the European Union’s Horizon Europe research and innovation programme under grant agreement No 101053962, Swedish State Support for Clinical Research (#ALFGBG-71320), the Alzheimer Drug Discovery Foundation (ADDF), USA (#201809-2016862), the AD Strategic Fund and the Alzheimer’s Association (#ADSF-21-831376-C, #ADSF-21-831381-C, and #ADSF-21-831377-C), the Bluefield Project, the Olav Thon Foundation, the Erling-Persson Family Foundation, Stiftelsen för Gamla Tjänarinnor, Hjärnfonden, Sweden (#FO2022-0270), the European Union’s Horizon 2020 research and innovation programme under the Marie Skłodowska-Curie grant agreement No 860197 (MIRIADE), the European Union Joint Programme – Neurodegenerative Disease Research (JPND2021-00694), the National Institute for Health and Care Research University College London Hospitals Biomedical Research Centre, and the UK Dementia Research Institute at UCL (UKDRI-1003). MZC is supported by the EU/European Federation of Pharmaceutical Industries and Associations Innovative Medicines Initiative 2 Joint Undertaking (IM2PACT grant no. 807015 and AIMS-2-TRIALS grant agreement no. 777394) and from National Institute for Health Research (NIHR) Oxford Biomedical Research Centre (BRC). A.K. received support through the National Institute for Health and Care Research.

