## Supplementary Methods and Figures for "iPSC-Astrocyte morphology reflects patient clinical markers"

### Supplementary Methods and Data

#### Differentiation of iPSC Astrocytes

Confluent iPSCs were cultured in 'Neural maintenance medium (NMM)' comprised of 50% Neurobasal (Life Technologies) and 50% DMEM-F12 media (Life Technologies) supplemented with 1% N2 (Life Technologies), 2% B27 (Life Technologies), 1% Glutamax (Life Technologies). This was supplemented with 10 µM SB-431542 (ApexBIO) and 1 µM LDN-193189 (Sigma) for 7 days with daily 100% media changes. Between days 7-10, cells were returned to neural maintenance medium. The day after, cells were maintained in NMM supplemented with 10ng/ml FGF-2 (Peprotech) and 10ng/ml EGF (Peprotech). Cells were maintained in this media and passaged when over 90% confluence as needed for 2-3 weeks at a density of 50,000 cells per $cm^2$. Cells were then switched to NMM supplemented with 10ng/ml EGF and 10ng/ml LIF (Peprotech) until astrocyte proliferation slowed at approximately 100 days. Astrocytes were matured for a further 50 days in NMM supplemented with 20ng/ml CNTF (Peprotech).

#### Harmony Pipeline

Harmony (PerkinElmer) software was used to analyse images captured by the Opera Phenix. Images were analysed for channel intensity (s100B, YKL-40 and NF-κB) to measure protein levels, general morphology measures including cell width-to-length. The pipeline included selecting first for nuclei, selecting a population where nucleus area was between >75$\mu m^2$ and <700$\mu m^2$, and nucleus roundness >0.75µm. Standard morphology properties were then calculated for cells in the s100B channel filtering cells out that were >240$\mu m^2$. Standard intensity properties were calculated for all channels.

#### Extended: Deep Learning: Classification Task and Similarity Learning

Images were classified using the s100B channel from all conditions and pre-processed by removing the brightest 0.4% pixels, normalised pixel intensity values and resized to 540*540. Images captured at three focal planes 0.5µm apart in the same field view were combined into a 540*540*3 image. The output of the model gave a vector of 16 values ranging from 0 to 1 that summed to 1, each of which represented the probability of the input image belonging to one of the 16 conditions. The and transfer learning applied, where the weights were pretrained on the ImageNet dataset [1]. The input layer of MobileNetV2 was replaced by our data structure (540*540*3). The classification layer was also removed, and a max pooling layer and global average pooling layer attached. The model adopted an end-to-end training manner, where it took a raw 3 layer z-stack image and directly yielded 16 numbers. The dataset was trained on 80% of the images and tested on the remaining 20%. Image augmentation techniques such as random flip, rotation and zoom, were applied to create transformed versions of images to artificially expand the dataset, which enabled the model to be more robust by introducing potential extra variations in the real world and prevent model overfitting.

To compare the similarity between each of the 16 conditions, the model was firstly trained on the previous training dataset and the weights/parameters of the model fixed. The images were then fed into the test dataset (15 images per condition) obtaining the output (a vector of 1,280 values) in the second to the last layer of our model, i.e., the global average pooling layer. Here, the vector of the 1,280 values were the highly extracted features of the input image, which can be viewed as a representation of the image. Cosine similarity was then used to evaluate and quantify the similarity between conditions. This measurement quantifies the similarity between two or more vectors and is a value that is bound by a constrained range of 0 and 1. Specifically, the cosine similarity is the cosine of the angle between vectors.

$$cosine\ similarity = \cos(\theta) = \frac{\boldsymbol{A} \cdot \boldsymbol{B}}{\|\boldsymbol{A}\|\|\boldsymbol{B}\|} = \frac{\sum_{i=1}^{1280} A_i B_i}{\sum_{i=1}^{1280} A_i^2 \sum_{i=1}^{1280} B_i^2},$$

where $\boldsymbol{A}$ and $\boldsymbol{B}$ are two different feature vectors. To calculate the image-level cosine similarity between two given conditions, one image from the first class was selected and the cosine similarity calculated between this image and all the images in the other condition, which yielded 15 values. We then looped this process through all the images in the first condition, thus yielding $15 \times 15 = 225$ values between the two conditions. The 225 values were then averaged to obtain a single value, representing the similarity between the two conditions. This was looped through all conditions and the similarity calculated for pairwise control and Aβ treatment conditions across cell lines.

**Quantitative PCR**

RNA was extracted using Trizol (Life Technologies) and cDNA synthesised using the TaqMan reverse transcriptase kit (Applied Biosystems) both according to manufacturer's instructions. cDNA was combined with FastStart Universal SYBR Green (Sigma-Aldrich) and the appropriate primers: *RPLP0* Forward 5'-GAAACTCTGCATTCTCGCTTCC Reverse 5'-GATGCAACAGTTGGGTAGCCA (housekeeping) and genes of interest *CHI3L1* (YKL-40) Forward 5'-GTGAAGGCGTCTCAAACAGG Reverse 5'-GGTCAAGGGCATCTGGGAAG and *S100B* Forward 5'-CAACAATGAGCTTTCCCAT Reverse 5'-CATTGTCCAGTGTTTCCATG**.** qPCR reactions were run using a StepOne Plus™ Real-Time PCR System (Applied Biosystems, 4376600). The cycle programme includes a denaturation step at 95°C for 3 minutes, followed by 40 cycles from 95°C for 20 seconds and 60°C for 30 seconds, heating from 60°C to 95°C for 15 seconds, and cooled to 60°C for 1 minute, and heated to 95°C for 10 seconds at increments of 0.5°C. Relative quantification of gene expression was analysed by the Pfaffl method [2].

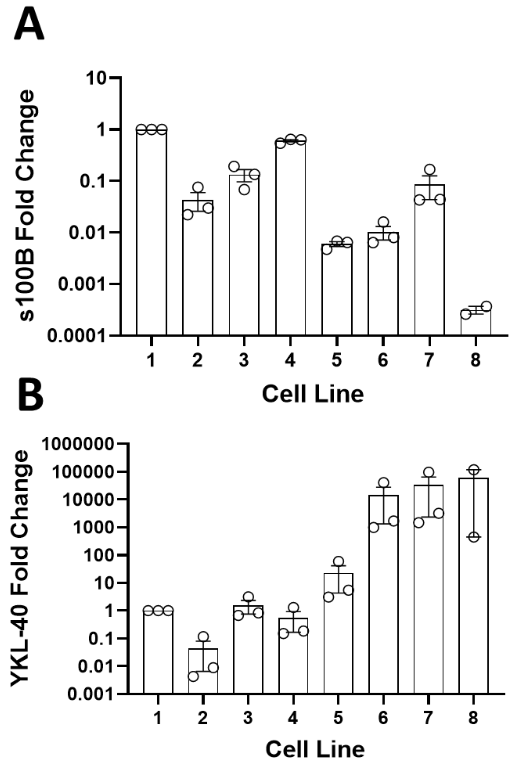


C

| Patient Number | Sex | APOE Genotype | CSF YKL-40 Concentration (pg/mL) |
|---|---|---|---|
| 1 | F | ε4/ε4 | 294426.012 |
| 2 | M | ε2/ε3 | 121037.573 |
| 3 | M | ε4/ε4 | 223639.5495 |
| 4 | F | ε3/ε4 | 164478.6165 |
| 5 | M | ε3/ε4 | 225878.8775 |
| 6 | M | ε3/ε3 | 137380.758 |
| 7 | F | ε4/ε4 | 246083.205 |
| 8 | M | ε3/ε4 | 133511.3575 |

**Supplementary Figure 1:** Fold change of (A) s100B and (B) YKL-40 at baseline between patients. s100B and YKL-40 mRNA is detected in all iPSC-astrocytes from patients but is variable. (C) Table demonstrates the APOE genotype and CSF YKL-40 concentrations from all patients used in this study. CSF YKL-40 levels classed as ‘high’ are shown in orange, ‘low’ levels are shown in green.
